# Competitive dewetting underlies site-specific binding of general anesthetics to GABA(A) receptors

**DOI:** 10.1101/694612

**Authors:** Sruthi Murlidaran, Jérôme Hénin, Grace Brannigan

## Abstract

GABA(A) receptors are pentameric ligand-gated ion channels playing a critical role in the modulation of neuronal excitability. These inhibitory receptors, gated by *γ*-aminobutyric acid (GABA), can be potentiated and even directly activated by intravenous and inhalational anesthetics. Intersubunit cavities in the transmembrane domain have been consistently identified as putative binding sites by numerous experiment and simulation results. Synaptic GABA(A) receptors are predominantly found in a 2*α*:2*β*:1*γ* stoichiometry, with four unique inter-subunit interfaces. Experimental and computational results have suggested a perplexing specificity, given that cavity-lining residues are highly conserved, and the functional effects of general anesthetics are only weakly sensitive to most mutations of cavity residues. Here we use Molecular Dynamics simulations and thermodynamically rigorous alchemical free energy perturbation (AFEP) techniques to calculate affinities of the intravenous anesthetic propofol and the inhaled anesthetic sevoflurane to all intersubunit sites in a heteromeric GABA(A) receptor. We find that the best predictor of general anesthetic affinity for the intersubunit cavity sites is water displacement: combinations of anesthetic and binding site that displace more water molecules have higher affinities than those that displace fewer. The amount of water displacement is, in turn, a function of size of the general anesthetic, successful competition of the general anesthetic with water for the few hydrogen bonding partners in the site, and inaccessibility of the site to lipid acyl chains. The latter explains the surprisingly low affinity of GAs for the *γ* − *α* intersubunit site, which is missing a bulky methionine residue at the cavity entrance and can be occupied by acyl chains in the unbound state. Simulations also identify sevoflurane binding sites in the *β* subunit centers and in the pore, but predict that these are lower affinity than the intersubunit sites.

**Significance:** After over a century of research, it is established that general anesthetics interact directly with hydrophobic cavities in proteins. We still do not know why not all small hydrophobic molecules can act as general anesthetics, or why not all hydrophobic cavities bind these molecules. General anesthetics can even select among homologous sites on one critical target, the GABA(A) heteropentamer, although the origins of selectivity are unknown. Here we used rigorous free energy calculations to find that binding affinity correlates with the number of released water molecules, which in turn depends upon the lipid content of the cavity without bound anesthetic. Results suggest a mechanism that reconciles lipid-centered and protein-centered theories, and which can directly inform design of new anesthetics.

## 1 Introduction

General anesthetics (GAs) are chemically diverse molecules that cause immobilization and amnesia with moderate potency. Their surprising structural diversity has motivated quantitative efforts to predict potency from molecular properties for over a century. The classic Meyer-Overton trend indicated that potency of GAs could be predicted by their solubility in olive oil (1), suggesting a lipid or membrane-mediated mechanism. It is now well established that direct binding of GAs to multiple proteins in the central nervous system, including ligand-gated ion channels (2–4), is essential to the effects of GAs at clinical concentrations. In particular, *γ*-aminobutyric acid receptors type A (GABA_A_), which are critical for rapid inhibitory signaling, have been identified as primary targets for widely-used general anesthetics like propofol and sevoflurane (5, 6).

GABA_A_ Rs are members of the pentameric ligand-gated ion channel (pLGIC) superfamily, and most synaptic GABA_A_R are heteropentamers with a 2*α*:2*β*:1*γ* stoichiometry. (7–9) Crystal structures of the *β*3 GABA_A_R homopentamer, (10) and recent cryo-electron microscopy structures of the *α*1*β*2*γ*2 heteropentamer (11, 12) and *α*1*β*3*γ*2 heteropentamer (13, 14) have confirmed that GABA_A_R share common structural features of the pLGIC family. Five subunits are arranged around a central pore; each subunit consists of an extracellular domain (ECD) containing at least two orthosteric ligand binding sites, a transmembrane domain (TMD) containing 4 helices notated M1-M4, with the M2 helix lining the pore and the M4 helix facing the lipid membrane, and a largely disordered intracellular domain.

General anesthetics (GAs) potentiate GABA_A_, but inhibit several other pLGICs, including some nAChRs and the prokaryotic channels GLIC and ELIC. Our initial computational studies on the nAChR predicted GA binding to multiple sites, including the pore lumen (15). Subsequently, the first crystal structure of GAs in complex with GLIC indicated binding in the the subunit center, and inhibition was attributed to this intrasubunit site (16). More quantitative simulations confirmed our pore binding hypothesis, and we argued that due to detergents in the crystallized GLIC pore, absence of crystallographic evidence for pore block by GAs was not decisive (17). Using Molecular Dynamics Simulation and Alchemical Free Energy Perturbation (AFEP) we predicted that, at half-saturating concentrations, propofol was as likely to bind to the GLIC pore as to GLIC intrasubunit sites, while isoflurane was significantly more likely to bind to the pore. Subsequent experimental results have increasingly supported a role for pore block in inhibition (18–20).

A pore binding site cannot, however, explain the gain of function that is observed in pLGICs responsible for the clinical effects of GAs on humans, and in the present manuscript we predict that propofol and sevoflurane have lower affinity for the pore binding sites of the GABA_A_ receptor than for several other potential allosteric sites. The only structures of GAs in complex with GABA_A_ (or any other eukaryotic receptor) come from co-crystals of GABA_A_ and several neurosteroids (21–23). These structures reveal a common binding site that is external to the TMD, at subunit interfaces. Based on the body of evidence from other co-crystal structures, photolabeling, mutageneis, and simulation, such a superficial site is not expected to explain the strong modulatory effects of GAs on GABA_A_. It is likely that the intrasubunit binding mode detected in GLIC causes gain of function, (20) as we originally suggested in (17), and possible that such an intrasubunit site could also cause gain of function in GABA_A_. However, numerous photoaffinity labelling studies have pointed to anesthetic binding sites at the interface between subunits in the GABA_A_ transmembrane domain (TMD). (24–30) This is illustrated in Figure 1, which represents data from (28, 29) using a propofol-based photolabel, and (30) using a sevoflurane-based photolabel. Perplexingly, mutation of most intersubunit residues (with a few exceptions) yields only subtle quantitative shifts in *EC*_50_(26, 31, 32) but affinity appears to be highly sensitive to subunit species. (29, 33, 34)

**Figure 1:**
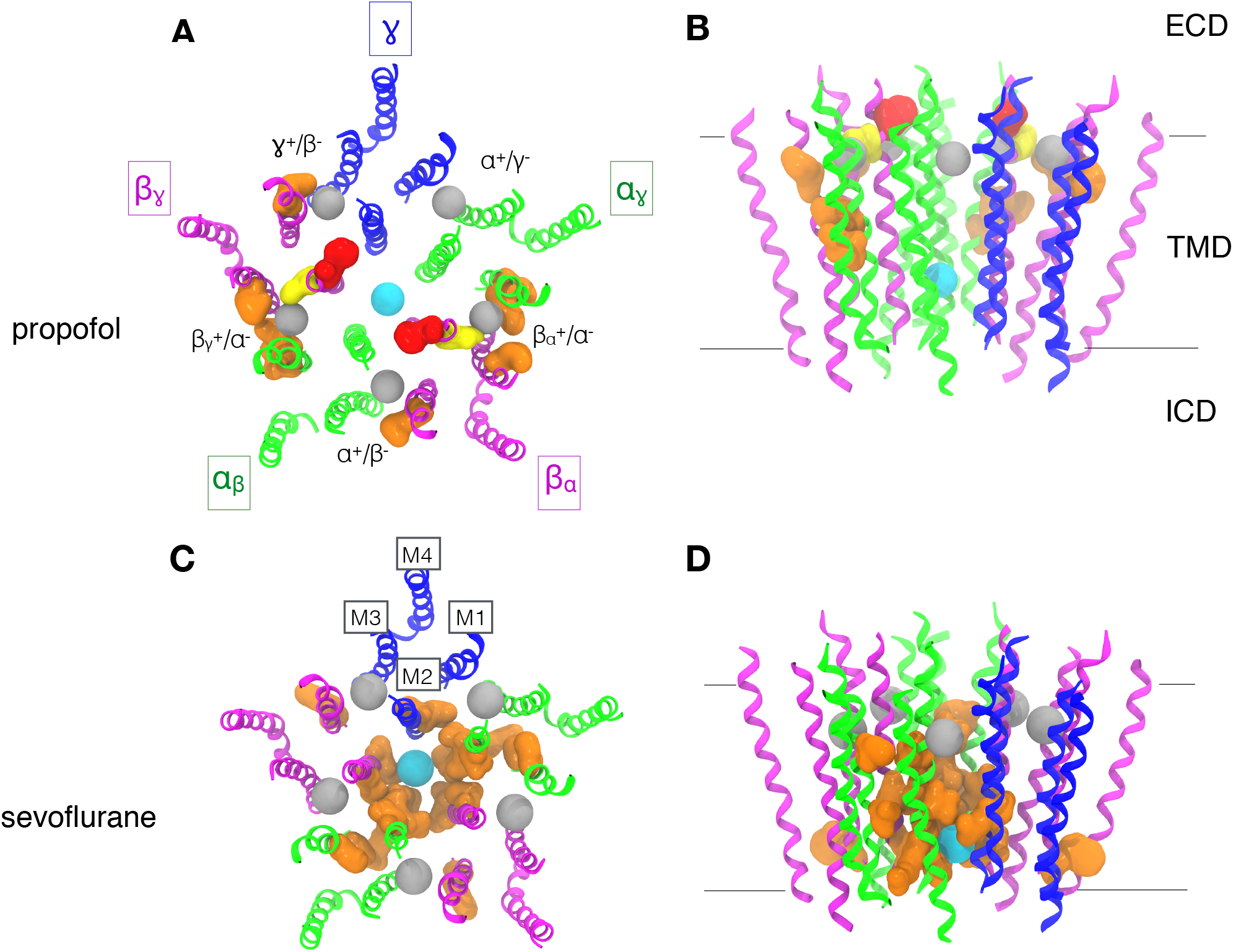
Experimentally-indicated contact residues for propofol and sevoflurane on the GABA_A_ receptor. The transmembrane domain of the GABA_A_R, viewed from the extracellular region (A,C) or from within the membrane (B,D), colored by subunit (*α*-green, *β*-purple,*γ*-blue). A-B) Propofol binding site residues indicated through photolabeling using AziPM(29) are shown in orange, o-PD(28) in red, and through mutagenesis(24) in yellow. The center of the intersubunit or pore binding cavities for propofol indicated by refinement simulations are marked with gray and cyan spheres, respectively. C-D) Sevoflurane binding site residues indicated through photolabeling with Azisevoflurane(30) are shown in orange. The center of the intersubunit or pore binding cavities for sevoflurane indicated by refinement simulations are marked with gray and cyan spheres, respectively. Both molecules were mobile in all intersubunit and pore sites during refinement simulation, and contact probabilities with individual residues are shown in Table 1.

Propofol has been a widely used general anesthetic since its discovery in 1980; it has a higher potency and is better tolerated than older, volatile anesthetics like halothane or isoflurane. Propofol potentiates GABA_A_ (26) at clinical (micromolar) concentrations and even directly activates the channel at higher concentrations (35, 36). In contrast, most volatile anesthetics potentiate GABA_A_ at 100 *μ*M concentrations (37).

The origins of the increased potency of propofol are not fully determined, despite studies involving a large number of propofol analogs, (26) but some evidence supports a role for the strong hydrogen bonding tendency of the hydroxyl moiety. A propofol analog with fluorine rather than hydroxyl (“fropofol”) has significantly reduced effects on GABA_A_ receptors, (38) and we previously reported (33) that propofol affinity for pseudosymmetric sites is an increasing function of hydrogen bonding propensity. One surprising result from (33) was the low affinity of propofol for the intersubunit site with the most polar residues (the *β*_*α*_^+^-*α*^−^ site). We report here that we previously underpredicted both hydrogen bonding and affinity for the *β*_*α*_^+^-*α*^−^ site, as a result of a technical artifact, specific to flexible docking of propofol and corrected in the present work.

Several observations are inconsistent with affinity driven by hydrogen bonding, however. Foremost, it is not clear why hydrogen bonding propensity would explain the strong correlation between potency and solubility in olive oil first observed by Meyer and Overton. Although isoflurane and sevoflurane do not contain a hydroxyl group like propofol, they do contain multiple weak hydrogen bond donors (39), and they are more soluble in water and less soluble in olive oil than propofol. While direct substitution of the propofol hydroxyl with fluorine yields an inactive molecule(38), substitution with bromide still leaves a general anesthetic as long as the solubility is maintained by modifying the isopropyl groups. (26) Furthermore, the *α*^+^-*γ*^−^ site within the GABA(A) receptor is a so-called “orphan site”(40) that has not been identified using any anesthetic photolabels, but contains the same number of hydrogen bonding residues(33) as the *α*^+^-*β*^−^ site that does bind azipropofol. (40)

In the present manuscript we use Molecular Dynamics Simulation and Alchemical Free Energy Perturbation to predict specific binding affinities across five subunit interfaces and two general anesthetics, and isolate the molecular origins of these affinity differences. We also find that water molecule displacement, rather than hydrogen bonding, is a more universal predictor of affinities for both sevoflurane and propofol across all five intersubunit sites. Water displacement, in turn, is determined by size of the anesthetic, lipid-accessibility of the intersubunit binding site, and the ability of the anesthetic to replace water for a limited number of hydrogen bonding partners in the binding site.

## 2 Methods

We use three general types of MD simulations for finding and ranking GA binding sites: (1) Docking refinement simulations, (2) Flooding simulations, and (3) Binding affinity calculations. Refinement and flooding simulations share the goal of generating poses and sampling interactions. Both use traditional equilibrium molecular dynamics approaches, but differ in the initial conditions and GA concentration. In refinement simulations, a small number of GA molecules begin in poses previously identified by docking, and may then reorient or even unbind during the simulation. In flooding simulations, a large number of GA molecules are placed in the water and spontaneously partition to the membrane and sites on the protein during the course of the simulation. Neither of these methods provides any quantitative information to rank sites, requiring that we calculate binding affinities using a different approach. We use the theoretically exact, alchemical free energy perturbation (AFEP) method to predict the GA concentration for which any specific site would be 50% occupied. Either refinement or flooding simulations can be used to provide initial coordinates for binding affinity calculations, but we primarily use refinement simulations. The workflow underlying our general approach for identifying and ranking binding sites is more explicitly detailed in Ref. (41).

### 2.1 Model systems for simulations

For straightforward comparison with previous work(33), we use a model of the *α*_1_*β*_3_*γ*_2_ GABA_A_ receptor based on the nematode GluCl crystal structure (3RHW) with ivermectin, a positive modulator, bound to the intersubunit sites in the TMD (42). This structure represents the active state, and the crystallized GluCl chain posesses 33 to 39% sequence identity with the GABA_A_ subunits studied here. Cryo-EM structures of human GABA_A_ receptor heteromers have been released recently(11–14), yet none of them is representative of the active state that is best suited to study binding of positive allosteric modulators. The systems were prepared as in references 43 and 33, by embedding the protein in a lipid bilayer composed of phosphatidylcholine (POPC), solvated and completed with 0.15 mol/L NaCl using CHARMM-GUI Membrane builder (44), with the final system containing 268 POPC molecules and about 180000 atoms.

### 2.2 Simulation details

All simulations used the CHARMM36 model (45) for proteins, phospholipids, ions, and cholesterol molecules, and the TIP3P water model (46). The forcefield parameters for sevoflurane were those used in (47), based on the isoflurane parameters developed in (39). Propofol parameters were those derived in (17), including the dihedral angle cross-terms (CMAP) required to adequately couple the hydroxyl and isopropyl groups of propofol. Energy minimization and MD simulations were conducted using the NAMD 2.12 package(48). All simulations employed periodic boundary conditions, long-ranged electrostatics were handled with a smooth particle mesh Ewald method, and a cutoff of 1.2 nm was used for Lennard-Jones potentials with a switching function starting at 1.0 nm. All simulations were run in the NPT ensemble with weak coupling to Langevin thermostat and a barostat at a respective 300 K and 1 atm. All bonds to hydrogen atoms were constrained using the SHAKE/RATTLE algorithm. A multiple time-step rRESPA method was used, with a high frequency time-step of 2 fs and a slow time-step of 4 fs for the long-range force components. All standard MD simulations started with 10,000 steps of energy minimization. Harmonic restraints with force constant 0.5 kcal/mol/Å^2^ on the protein backbone positions prevented conformational drift during the AFEP calculations.

### 2.3 Docking and MD refinement

General anesthetics were docked to intersubunit sites using the Autodock Vina software (49). The search space comprised the entire TM domain and intersubunit sites. Poses in each of the five intersubunit sites as well as the pore were chosen after performing multiple docking runs as described in Ref. (41). Although propofol bonds were rotatable during docking for our previous studies(17, 33), here they were made non-rotatable, and only the low-energy conformation of propofol was used for docking, as justified in Ref. (41).

Unbiased docking refinement simulations involving propofol were run for about 600 ns following minimization. Propofol remained stable in the binding site throughout the course of the simulation. Autodock Vina initially docked sevoflurane to sites in the lower TMD, but during MD refinement for about 200 ns, sevoflurane reliably shifted higher in the TMD in three of the intersubunit sites, to a position more similar to that of propofol (41). Consequently, all initial sevoflurane configurations here use the “upper” site. In line with the faster dynamics of sevoflurane compared to propofol, these MD simulations of sevoflurane were run for about 150 ns, with sevoflurane remaining stably bound despite being highly mobile in the site.

### 2.4 Free energy perturbation simulations

Alchemical free energy perturbation simulations were performed on general-anesthetic bound GABA_A_R systems, with starting configurations obtained from standard MD simulations. Movement of the ligand was confined to 5 Å of the binding site center by flat-bottom restraints, which were spherical in the intersubunit sites and cylindrical in the pore, with a force constant of 5 kcal/mol/Å^2^. Decoupling of the ligand was performed over a series of 24 windows. The perturbation parameter λ was sampled with a step size equal to 0.025 between 0 < λ < 0.1 and 0.9 < λ < 1, and a step of 0.05 otherwise. Each window began with a 4 ps equilibration period followed by a 5 ns run for data collection. Lennard-Jones interactions at intermediate λ values were interpolated as a smooth, “separation-shifted” soft-core potential (50) as implemented in NAMD. Free energy of binding was calculated following Ref. (17), as:

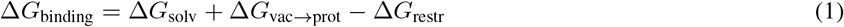

where ΔG_solv_ is the solvation free energy of the anesthetic, ΔG_vac→prot_ prot is the free energy decoupling the ligand from the binding site and ΔG_restr_ if the free energy cost of imposing restraints relative to the standard state volume. As these calculations deal with relatively small ligands with significant orientational mobility in the binding site, simple center-of-mass restraints are sufficient to ensure rapid convergence. This would not be the case for larger ligands with slow orientational and conformational fluctuations, which would benefit from more specialized restraints.(51)

### 2.5 Flooding simulations

To identify additional candidate sites, we used an approach similar to that used for isoflurane in Ref. (15): sevoflurane was inserted randomly into the water surrounding the protein using tcl scripts within VMD (52), with a sevoflurane-to-lipid ratio of about 1:3. Flooding simulations were run for about 1.7 *μ*s. Flooding with propofol was not pursued because the extremely low solubility causes aggregation of ligand in aqueous solution; sufficiently low concentrations prevent aggregation but extend the required time scales beyond those feasible for simulation. As discussed in Ref. (41), this approach is not straightforward for propofol, due to its low solubility.

### 2.6 Analysis

All molecular graphics was created using VMD (52). In flooding simulations, sevoflurane was determined to have partitioned to the membrane if there are at least 3 acyl chains within 5 Å of sevoflurane, in the protein if there are at least 3 protein residues within the 3 Å of sevoflurane, and in the aqueous phase otherwise. These criteria were chosen so that the sevoflurane partitioning status consistently matched visual inspection. Solvation of the binding cavities was determined by counting the number of lipid atoms or water molecules that met the following criteria per frame: coordinates within 5 Å from the center of the binding cavity. Hydrogen bonds were determined using geometric criteria, with a distance cutoff of 3.3 Å length and angle cutoff of 40 degrees. Solvation and hydrogen bonds were calculated for every ns, and the average was calculated after discarding the first 50 ns of simulations.

## 3 Results

### 3.1 Simulations show frequent contacts between general anesthetics and residues identified by photolabeling

Various residues in the intersubunit sites of GABA_A_R have been identified using photoaffinity labeling as possible binding sites for propofol (29) and sevoflurane (30), as shown in Figure 1. For comparison, the contact frequency and mean distance measured from simulations are listed in Table 1. These values are extracted from refinement simulations following docking to the interface or pore sites.

**Table 1:**
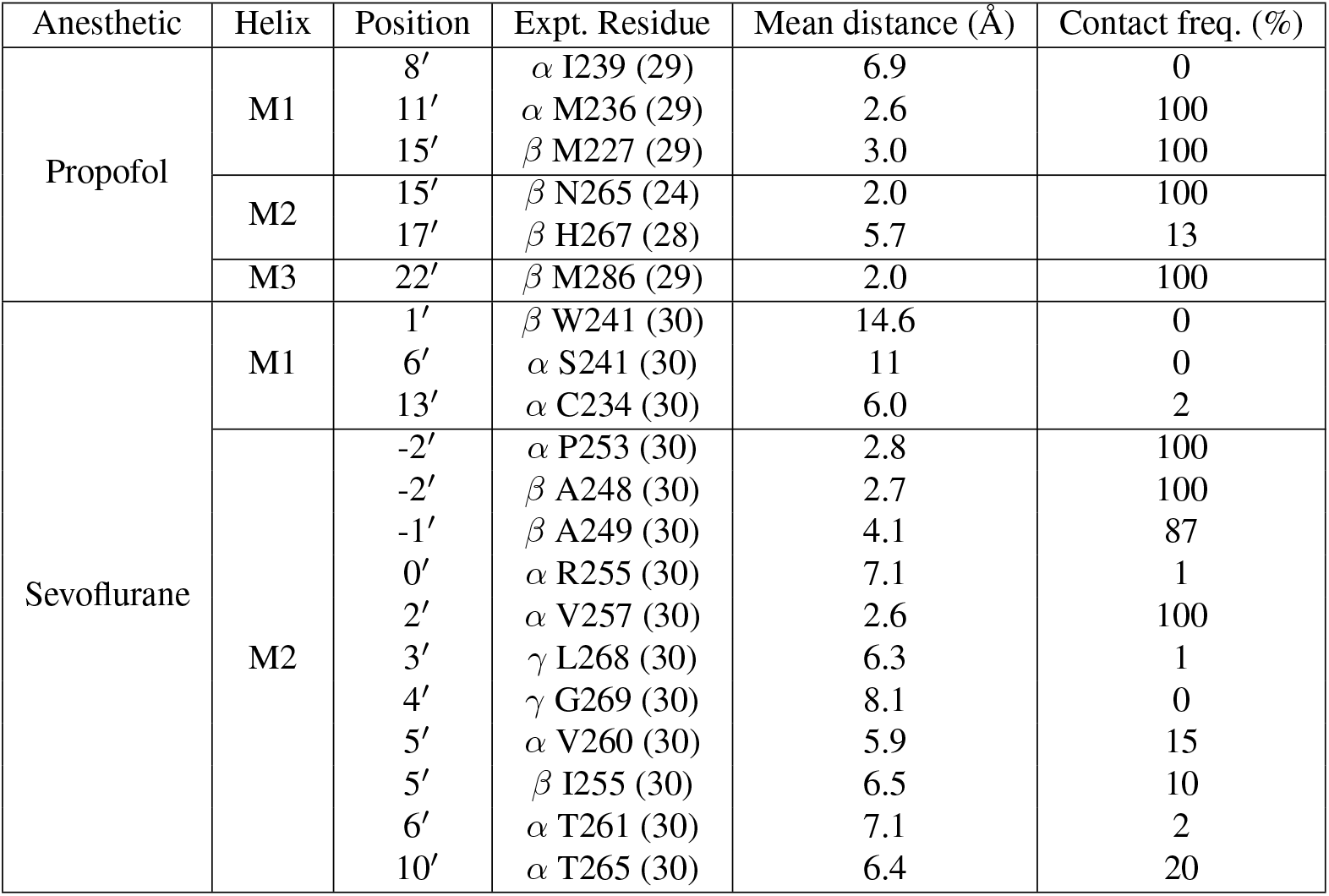
Contacts between general anesthetics and experimentally-identified contact residues, sampled during refinement simulations in which a single general anesthetic molecule occupied all intersubunit cavities and the pore. Experimentally-identified residues are also shown in Figure 1. Contacts are characterized by both the mean distance throughout the simulation and the percentage of sampled frames in which the distance was below a 5 Å cutoff.

| Anesthetic | Helix | Position | Expt. Residue | Mean distance (Å) | Contact freq. (%) |
| --- | --- | --- | --- | --- | --- |
| Propofol | M1 | 8' | $\alpha$ I239 (29) | 6.9 | 0 |
| | | 11' | $\alpha$ M236 (29) | 2.6 | 100 |
| | | 15' | $\beta$ M227 (29) | 3.0 | 100 |
| | M2 | 15' | $\beta$ N265 (24) | 2.0 | 100 |
| | | 17' | $\beta$ H267 (28) | 5.7 | 13 |
| | M3 | 22' | $\beta$ M286 (29) | 2.0 | 100 |
| Sevoflurane | M1 | 1' | $\beta$ W241 (30) | 14.6 | 0 |
| | | 6' | $\alpha$ S241 (30) | 11 | 0 |
| | | 13' | $\alpha$ C234 (30) | 6.0 | 2 |
| | M2 | -2' | $\alpha$ P253 (30) | 2.8 | 100 |
| | | -2' | $\beta$ A248 (30) | 2.7 | 100 |
| | | -1' | $\beta$ A249 (30) | 4.1 | 87 |
| | | 0' | $\alpha$ R255 (30) | 7.1 | 1 |
| | | 2' | $\alpha$ V257 (30) | 2.6 | 100 |
| | | 3' | $\gamma$ L268 (30) | 6.3 | 1 |
| | | 4' | $\gamma$ G269 (30) | 8.1 | 0 |
| | | 5' | $\alpha$ V260 (30) | 5.9 | 15 |
| | | 5' | $\beta$ I255 (30) | 6.5 | 10 |
| | | 6' | $\alpha$ T261 (30) | 7.1 | 2 |
| | | 10' | $\alpha$ T265 (30) | 6.4 | 20 |

Propofol formed frequent contacts with three residues labeled by the photolabel azipropofol (29) and one residue indicated by mutagenesis (24), which line cavities between all subunit interfaces except *α*^+^-*γ*^−^ (Figure S1). These residues are all located near the bulge in M1 at 18′ and 19′. Two residues located farther from the bulge were not frequently contacted. One such residue (*β* M2 H267) was labeled by ortho-azipropofol (28), which does not compete with propofol. The other residue (*α* M1 I239) is on the intracellular end of the same interface as two residues (*α* M236 and *β* M286) which we propose play a dominant role in forming a high-affinity site.

Residues labeled by azisevofluorane (30) are concentrated in the intracellular half of the TMD (Figure 1D). They are primarily pore-lining residues on the M2 helix, with some on M1 facing the lipid membrane or the intrasubunit cavity. Since there are no residues labeled on M3, it is not possible to distinguish intersubunit vs intrasubunit binding modes, on the basis of residues labeled in Reference (30) alone. Interpretation of the photolabeling results in (30) relied heavily on complementary docking calculations based on the present structural model of the *α*_1_*β*_3_*γ*_2_ GABA(A) receptor. As discussed in Ref. (41), docking with volatile anesthetics is especially prone to false positives, and poses identified by docking may be found to be unstable or short-lived. To avoid this pitfall, we refined docking poses for sevoflurane docked to both intersubunit and pore sites using molecular dynamics simulation. The frequencies of sevoflurane contacts with photolabeled residues after 200 ns of docking refinement are shown in Table 1. In these refinement simulations, we rarely observed contacts between sevoflurane and the M1 photolabeled residues. If sevoflurane bound to an intersubunit site has little access to these M1 residues, the experimental results could instead be explained by the photolabel binding to the *α* intrasubunit site rather than an intersubunit site. Most of the labeled residues in Reference (30) are on the M2 helix, and many of those face the pore in both the homology model and a very recent EM structure (14). In our view, these residues collectively suggest heavy labeling of the pore. Sevoflurane and the photolabel used in Ref. (30) both potentiate GABA(A) receptors, which is not consistent with pore block, but sevoflurane is a small, mobile molecule that may occupy the pore transiently without blocking it. We discuss the particular case of pore-binding in a subsequent section.

### 3.2 Sevoflurane spontaneously binds to intrasubunit, intersubunit, and pore sites

Flooding simulations involve introducing a high concentration of the ligand into the system (usually the water) and waiting for it to spontaneously partition into the membrane and/or bind to protein cavities. This approach can provide qualitative insight, by confirming that a reasonable pathway exists for putative sites, indicate sites where multiple occupancy should be considered, and suggest new putative sites. (15) Due to the limited number of observed binding and unbinding events, it is suitable for hypothesis generation but not quantitative calculations of binding affinities. Furthermore, flooding with molecules with very low solubility (such as propofol) is prohibitive, due to the high concentrations of ligand required. In the past, this approach has allowed us to accurately predict binding sites of isoflurane to the nAChR and GLIC, (15) while here we apply it to sevoflurane.

The fraction of sevoflurane partitioning into the membrane, water, or protein sites was tracked throughout the flooding simulation (Figure 2 A,B) and all sevoflurane molecules partitioned into membrane or protein sites by the simulation conclusion. Occupancy of protein sites was further broken down into intrasubunit, pore, and intersubunit, with ligand densities averaged over the final 1200 ns shown in Figure 2D. In addition to expected partitioning to intersubunit sites, sevoflurane was observed partitioning to the intrasubunit cavity of both *β* subunits (but not *α* or *γ* subunits), consistent with the binding of propofol and desflurane observed in GLIC co-crystals. (16) In both *β* subunits, sevoflurane binds between the M1 and M4 helix, while a lipid acyl chain enters the subunit center from the M3 and M4 helix. Unlike the *α* and *γ* subunits, the GABA(A)r *β* subunit has a large cavity in the center that is free of protein density, similar to what we observed previously for multiple nAChR subtypes.(53, 54)

**Figure 2:**
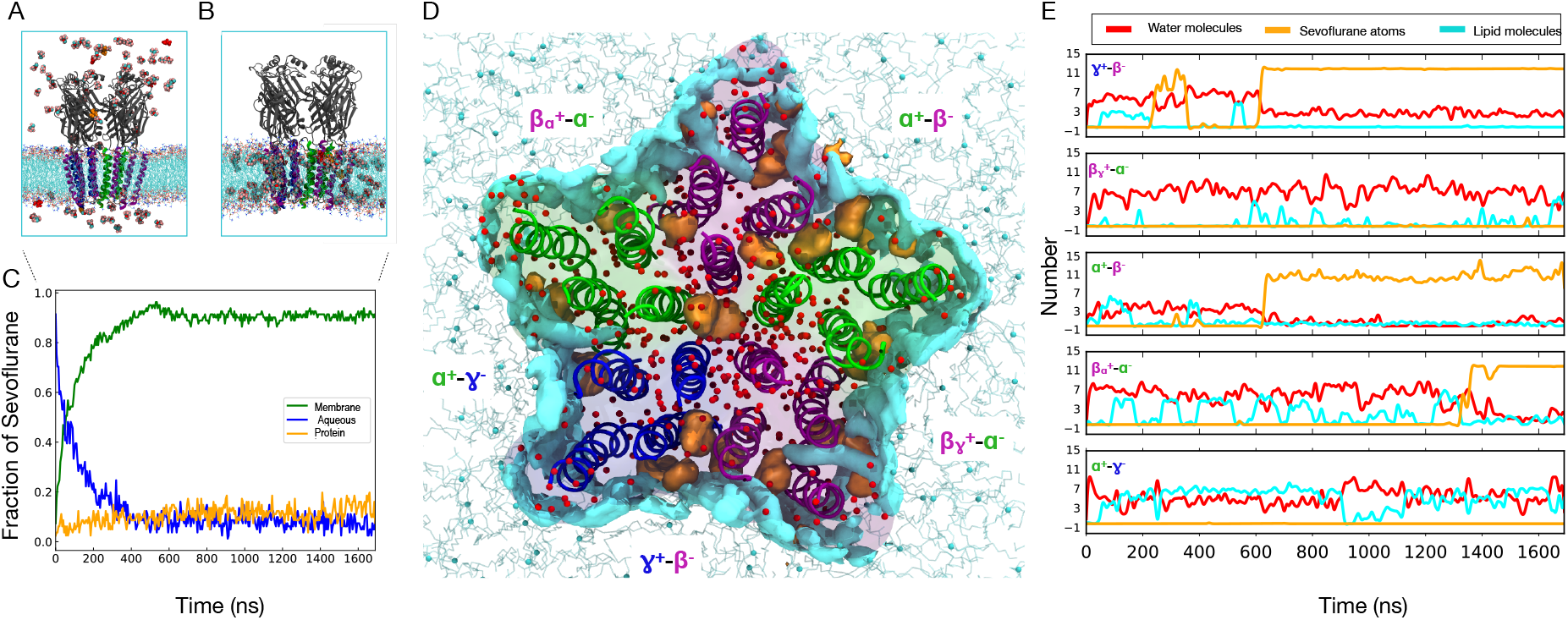
Flooding simulation of GABA_A_R with sevoflurane. (A) GABA_A_R embedded in lipid membrane, flooded with sevoflurane (colored by element) with the outline of the simulation box. (B) GABA_A_R system after the sevoflurane partitions into the lipid membrane. (C) Fraction of sevoflurane molecules in membrane, protein and aqueous phase. (D) View of the TMD of GABA_A_R from the ECD at the final frame of the flooding simulation, displaying water molecules (red), average density of sevoflurane molecules (orange) and lipids atoms (cyan) bound to the TMD of the channel. (E) Number of water (red) molecules, lipid atoms (cyan) and sevoflurane (orange) atoms that enter each intersubunit cavity in the course of the simulation.

Three sevoflurane molecules enter the pore through the ECD vestibule over 1.3 *μ*s, consistent with multiple occupancy of the pore observed in flooding/equilibrium simulations of GLIC and nAChR exposed to isoflurane, (15, 55) as well as later crystal structures. (19, 20) Since one of the three pore-bound sevoflurane molecules spontaneously shifted from the pore to the *α*^+^-*β*^−^ inter-subunit site, the pore occupancy was at most two sevoflurane molecules. Unlike GLIC, ELIC, or nAChR, GABA_A_ receptor function is enhanced by sevoflurane, and this result is only consistent with functional results if sevoflurane has lower affinity for the pore than for other TMD binding sites. This is confirmed via free energy calculations in Table 2.

**Table 2:**
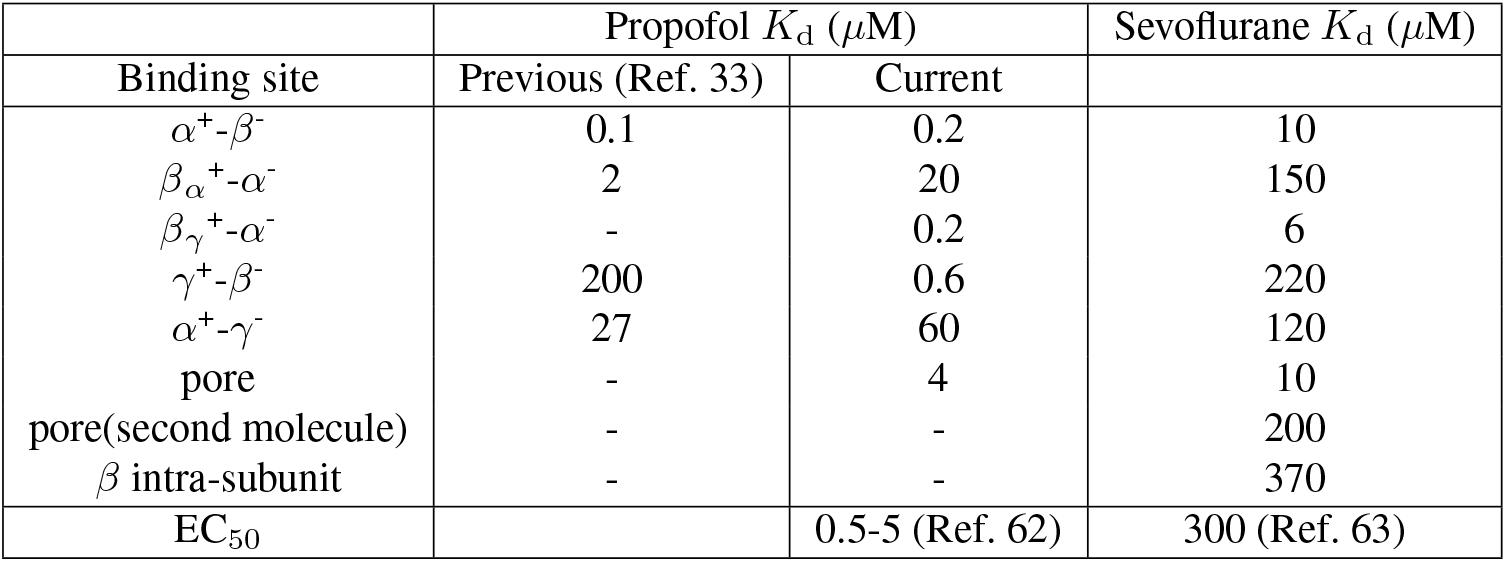
Binding affinities of propofol and sevoflurane for multiple sites on the GABA_A_ receptor, calculated using Alchemical free energy perturbation, including comparison to previous values we reported in (33). We discuss the substantial change in calculated affinity for propofol to the *γ*^+^-*β*^−^ site in the text. EC_50s_ for both propofol and sevoflurane reflect enhancement of GABA-activated chloride current in neurons incorporating several receptor subtypes. They indicate an upper bound on reasonable *K_D_* values, rather than an expected equality.

Visual observation of sevoflurane molecules at the conclusion of the flooding simulations reveals multiply occupied cavities, particularly the *α*^+^-*β*^−^ intersubunit cavity and the pore. (Figure 2E and F). In other inter and intrasubunit cavities, sevoflurane consistently interacts with lipids, which occupy *α*^+^-*γ*^−^ throughout the simulation and transiently bind *β_γ_*^+^-*α*^−^ site as well. Multiple occupancy of one binding cavity can not be feasibly identified via photoaffinity labeling or mutagenesis.

### 3.3 Both Propofol and sevoflurane form hydrogen bonds with the M1 backbone of γ and β, but not α

The fraction of frames in which the GA forms hydrogen bonds with each residue in the intersubunit cavities was calculated for the refinement MD simulations (Figure 3).

**Figure 3:**
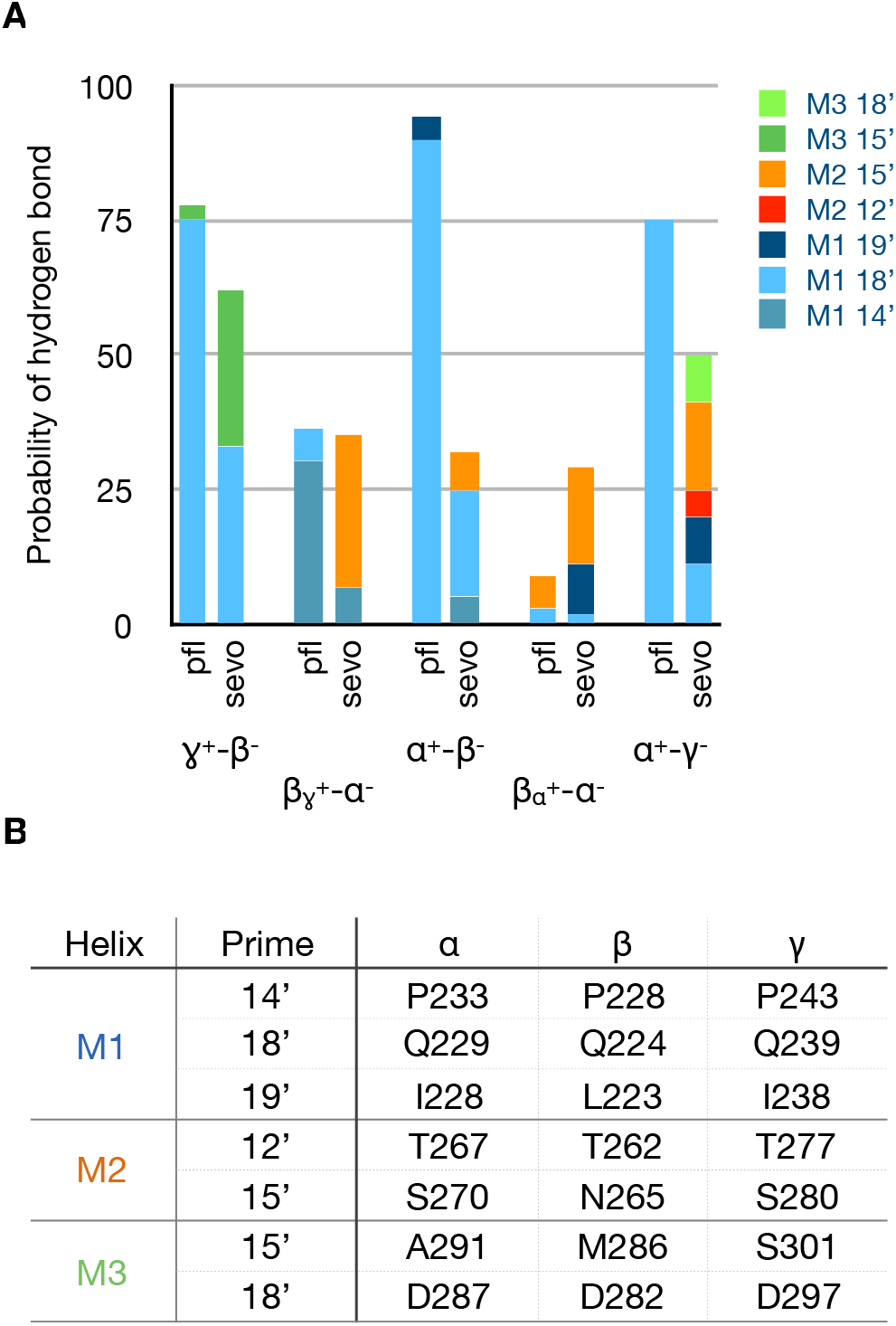
Frequency and distribution of hydrogen-bonding interactions. A) Fraction of refinement simulation frames in which a hydrogen-bond was formed between an intersubunit-bound GA and different residues facing the cavity. Each bar represents a different intersubunit cavity and anesthetic, while the hydrogen-bonding partner residues are represented by stacks within the bar. Hydrogen bonds formed with M1 18′ and M1 19′ involve the backbone carbonyl oxygen. B) Prime numbering index for residues listed in (A), following (56).

The conserved proline at M1 14’ (56) creates a bulge in the helix at the 18’ and 19’ positions, freeing the carbonyl oxygens at those positions for hydrogen bonding with an intersubunit ligand (specific residues are shown in Figure 3). Similar to the results of our previous work (30, 57), we observed direct hydrogen bonding of the intersubunit-bound ligand with M1 18′ and 19′. Both propofol and sevoflurane form such hydrogen bonds with this bulge in either *β* or *γ* M1 helices (i.e. in interfaces where the – face is either *β* or *γ*). This is consistent with the 62% sequence identity between the *β*_3_ and *γ*_2_ M1 helices.

In contrast, hydrogen bonding with the bulge on *α* subunits in the *β*+/*α*-sites was infrequent, for both anesthetics. Propofol was more likely to form hydrogen bonds either 1 turn lower, at *α*P238 (M1 14′) or reorient and form hydrogen bonds with *β* N265 (M2 15′) (Figure S3). Sevoflurane was also more likely to hydrogen bond with *β* N265 than with other residues in the cavity (Figure S3). M2 15′ has been the subject of multiple mutagenesis studies for numerous general anesthetics(6, 27, 58–60) and an extensive study on propofol suggesting a role for both affinity and efficacy. (61). The present results are consistent with favorable interactions between general anesthetics and polar sidechains at M2 15′, but suggest that hydrogen bonding with the backbone could compensate if these interactions were lost due to mutation.

As shown in figure 3, although sevoflurane is a weak hydrogen bond donor and does hydrogen bond less frequently overall than propofol within a given site, it has more hydrogen bonding partners. The large number of weak hydrogen-bonding partners is consistent with the presence of multiple donors in the ligand and high mobility of sevoflurane in the binding cavity.

### 3.4 Full occupancy of the GABA(A) receptor pore requires high concentrations of general anesthetics

Converging qualitative evidence indicates binding of anesthetics to the pore: previous photolabeling with azisevoflurane indicated a high number of pore-lining residues (Figure 1C and D), docking calculations report a GABA(A) receptor pore binding site for both propofol and sevoflurane, and sevoflurane spontaneously binds to the pore in flooding simulations (Figure 2). Yet, both molecules cause gain-of-function in GABA(A) receptors at low concentrations, indicating that the affinity for the pore must be lower than for the sites causing gain of function. We used AFEP to calculate affinities of sevoflurane and propofol for the pore as well as the intersubunit transmembrane binding sites (Table 2). For propofol, we calculate a *K_D_* for the pore of 4 *μ*M, which is weaker than three of the intersubunit sites, and a greater concentration than the typical *EC*_50_ of 0.5-5 *μ*M (62) This result is a useful sanity check for our approach, and is consistent with gain-of-function caused by binding to one or more of these three sites (*α*^+^-*β*^−^, *β*_*α*_^+^-*α*^−^ and *β*_*γ*_^+^-*α*^−^).

For particularly small molecules binding to the pore (including volatile anesthetics), it is critical to consider the possibility of multiple occupancy, for two reasons: 1) The binding site is so much larger than the ligand that multiple occupancy is not sterically unfavorable. In flooding simulations we observe double occupancy of the pore by sevoflurane, and the previously photolabeled residues line a much longer region of the channel lumen than a single sevoflurane molecule would contact. 2) Such molecules are so small and fluid in the pore that a single such molecule may not block conduction. We presented a functional model accounting for a mixture of partially and fully blocked pores in Ref. (17), where the probability that a single molecule would block conduction was a tunable parameter. For the affinities we measured to be consistent with the experimentally measured *IC*_50_, a single isoflurane molecule bound to the GLIC pore would block about 12% of the time.

We calculated affinities for a single sevoflurane molecule binding to the pore, and a second molecule binding to a pore that was already occupied (Table 2). The single sevoflurane molecule had an affinity of 10 *μ*M, comparable to the strongest intersubunit affinities and below the *EC*_50_ of 300 *μ*M (63). The second sevoflurane molecule, however, had a significantly lower affinity of 200 *μ*M, which is consistent with the pore unbinding incident we observed in flooding simulations. This negative cooperativity is consistent with what we measured for isoflurane binding to the closed GLIC pore (2.8 *μ* M for the first molecule, vs 80 *μ* for the second). In that case we also calculated a much lower affinity (2,800 *μ*M) for the putative allosteric site, suggesting that GLIC is inhibited by general anesthetics because pore block requires a much lower concentration than occupancy of a gain-of-function site. Our results here suggest the reverse trend for GABA(A)r, consistent with gain-of-function observed at low concentrations.

### 3.5 Methionine residues determine lipid accessibility to intersubunit cavities

In the apo state, both water molecules and lipid acyl chains occupy the intersubunit cavities, with the relative distribution depending on the cavity sequence (Figure S4). The average number of water molecules and lipid atoms per intersubunit site was determined for the apo and bound configurations (Figure 4A), and also tracked over the course of the AFEP simulations.

**Figure 4:**
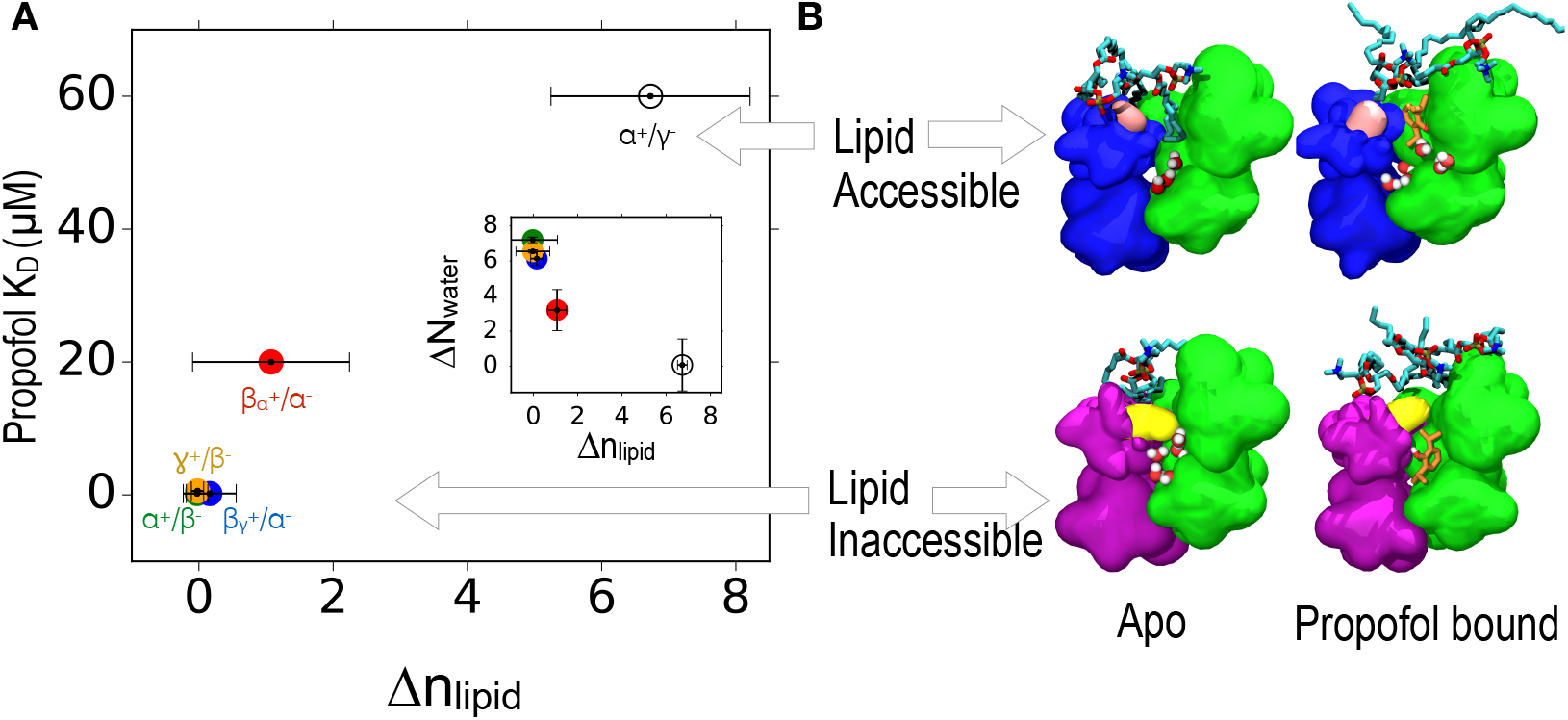
Solvation of lipid accessible and lipid inaccessible sites. (A) Correlation between average number of lipid molecules displaced in each intersubunit site: *γ*^+^-*β*^−^ (orange), *β*_*γ*_^+^-*α*^−^ (blue), *α*^+^-*β*^−^ (green), *β*_*α*_^+^-*α*^−^ (red), and *α*^+^-*γ*^−^ (white), and binding affinity of that site for propofol. Black error bars give the standard deviation for each site. Inset plot (in A) shows the correlation between average number of displaced water molecules (Δ*N_water_*) and average number of displaced lipid atoms (Δ*n_lipid_*) in each intersubunit site. (B) Snapshots of the intersubunit sites showing representative frames of the lipid accessible and inaccessible cavities in apo and propofol-bound receptor (propofol colored orange), viewed from the ECD along the pore axis. The *β*M227 (M1-15′) methionine residue forming the lipid-inaccessible cavity is colored in yellow; the homologous *γ* I242 (M1-15′) isoleucine residue forming the lipid-accessible cavity is colored in pink.

We find that in the apo state, the sites containing *α* and *β* subunits have a similar number of water molecules, while the *γ*^+^-*β*^−^ site has slightly more due to an additional polar residue, as shown in our previous study (33). The number of water molecules decreases with binding for most sites, consistent with the observations of Tang and co-workers.(64)

We also observe that the number of lipid atoms occupying the *α*^+^-*γ*^−^ site is significantly larger than for other sites. As depicted in figure 4 B, a bulky M286 at M3:22’ *β* in sites *β*_*α*_^+^-*α*^−^ and *β*_*γ*_^+^-*α*^−^ and M227 at M1:15’ *β* block lipid access for all intersubunit cavities but the *α*^+^-*γ*^−^ interface: we refer to these cavities as “lipid-inaccessible”. The *α*^+^-*γ*^−^ interface contains an M1 helix from *γ* subunit, with the M1:15’ Met replaced with a slightly less bulky Ile. Acyl chains of the surrounding phospholipid membrane can more readily access this site, and thus we term it a “lipid-accessible” cavity. Other intersubunits sites have a methionine residue at the 15’ or 22’ position in either the M1 of a subunit or the M3 of the neighboring subunit forming the interface. As shown in the inset, lipid displacement by propofol is inversely correlated to water displacement; below, we show that water displacement is the best predictor of binding affinity for both sevoflurane and propofol. Displacement of water molecules by the ligand, however requires that water molecules (not lipids) occupy the cavity in the apo state.

### 3.6 Affinity directly correlates with water displacement upon binding

In Ref. (33), we found a direct correlation between the likelihood of hydrogen bonding and affinity of general anesthetics for binding sites. One surprising outcome was the low hydrogen bonding propensity and low affinity of propofol for the *γ*^+^-*β*^−^ site, despite a large number of hydrogen bonding partners. We now believe that those results were artefactual outcomes of designating propofol’s isopropyl groups as rotatable during the docking calculations. We have resolved this technical issue as described in Methods, and we present updated affinities in Table 2. We now predict that *γ*^+^-*β*^−^ will be among the highest affinity cavities, whereas our previous calculations ranked it as the lowest affinity intersubunit cavity, with an affinity of 200 *μ*M.

Other affinities remain similar to their previous values, which reflects the ability of the propofol to relax rapidly from the unfavorable conformation in all but the tightest binding sites. A second unexplained outcome in Ref. (33) was the low affinity and low hydrogen bonding propensity of propofol for the *α*^+^-*γ*^−^ site, which had the same number of hydrogen bonding partners as the higher affinity *α*^+^-*β*^−^ site. In the present study, we find that propofol in the *α*^+^-*γ*^−^ site does form hydrogen bonds about as often as in the *α*^+^-*β*^−^ site (Figure 3) but still measure a particularly low affinity for the *α*^+^-*γ*^−^ site. In general for lipid-inaccessible sites, *K_D_* for propofol does tend to decrease with increased hydrogen bonding, as we observed in Ref (33), but this trend does not extend to the lipid-accessible *α*^+^-*γ*^−^ site, or for sevoflurane.

Given the unique lipid accessibility of *α*^+^-*γ*^−^, we considered whether affinity was more directly correlated to displacement of water or lipid. As shown in Figure 5 A, propofol *K_D_* decreases monotonically with the number of displaced water molecules and increases monotonically with the number of displaced lipid atoms (Figure S5, row 1). This suggests that the intersubunit cavities generally prefer to be occupied by hydrophobic groups rather than water. For lipid-inaccessible cavities, occupation by lipids is obviously not an option, but they can be occupied by small hydrophobic molecules like propofol. In all high affinity (*K_D_* < 1*μ*M) intersubunit propofol sites, more than five water molecules are displaced on average upon ligand binding.

**Figure 5:**
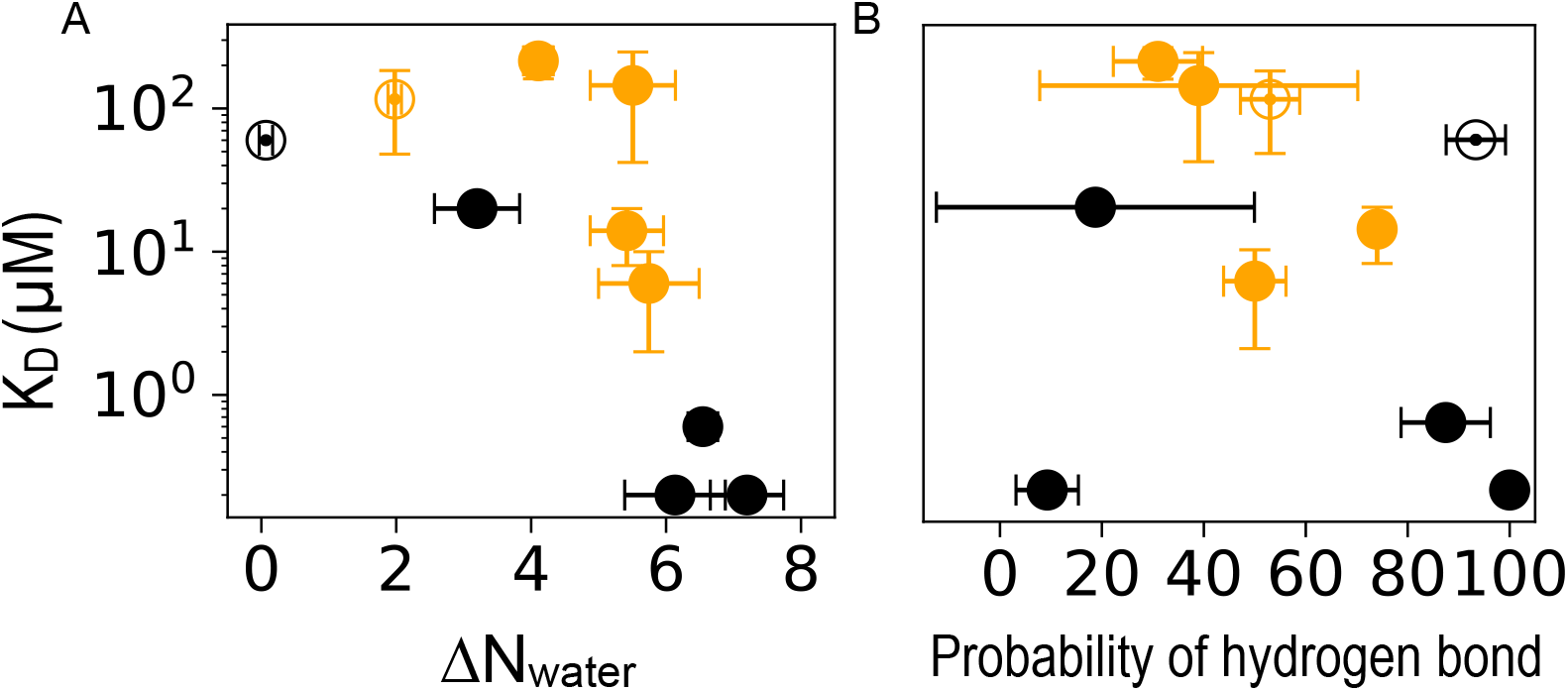
Intersubunit binding affinity as a function of displaced water atoms and hydrogen bonding. Correlation between binding site affinity (on a log scale) and (left) the average number of displaced water molecules *N_water_* or (right) the probability of a hydrogen bond being formed between propofol (black) or sevoflurane (orange) and a side-chain residue. The open and closed circles represent sites that are lipid-accessible and lipid-inaccessible respectively, as illustrated in Figure 4.

Very recent photolabeling protection studies(34) have indicated that propofol has a five-fold higher affinity for the two *β*^+^ sites (*β*_*α*_^+^-*α*^−^ and *β*_*γ*_^+^-*α*^−^) than for the two *β*^−^ sites (*α*^+^-*β*^−^ and *γ*^+^-*β*^−^). As shown in Table 2, we have measured a range of values (0.2 – 20 *μ* M) across three calculations involving *α*^+^/*β*^−^ sites. Interestingly, we find that for individual cavities, the variation in affinity from the mean also correlates with the variation in water displacement across replicas. As shown in Figure 4, an average of six water molecules is displaced from the *β_γ_*^+^-*α*^−^ site, while an average of three water molecules is displaced from the *β_α_*^+^-*α*^−^ site; the calculated affinities for the two sites are 0.2 *μ*M and 20 *μ* M respectively. Differences in the initial number of lipid atoms and water molecules across two sites with identical sequences can result from the initial lipid bilayer packing process, as well as from thermal effects; we can not interpret these results as indicating an intrinsic difference based on location within the pentamer. This observation suggests that endpoint solvation can serve as a primary source of statistical noise in AFEP calculations of membrane proteins.

Results for sevoflurane are consistent with water displacement as a determinant of affinity (Figure 5 A). Sevoflurane is a smaller molecule than propofol and does not displace more than five water molecules from any site, but the two sites with the highest number of displaced water molecules are still the two sites with the strongest affinity (Figure S5, row 2). Observing clearly monotonic trends across the sevoflurane data is challenging due to the large uncertainty in affinities and number of displaced atoms for the lowest affinity sites.

Remarkably, however, we find that sevoflurane and propofol affinity data exhibit the same trend with respect to either the number of water molecules or lipid atoms displaced, but not the number of hydrogen bonds formed (Figure 5). This suggests that the increased potency of propofol relative to sevoflurane may be primarily due to its larger molecular volume, and the resulting increased tendency to displace water from the cavity.

## 4 Conclusion

We predict the isolated binding affinities for sevoflurane and propofol of five transmembrane intersubunit cavities and the pore of a heteromeric GABA_A_ receptor, using MD simulation and the AFEP method. Our results exhibit mechanisms that answer two significant questions regarding the interactions of general anesthetics and GABA_A_ receptors.

The first longstanding question concerns the fundamental molecular interactions determining general anesthetic activity, and in particular, potency at the GABA_A_ receptor. We propose here that dewetting of hydrophobic cavities upon anesthetic binding is the primary determinant of affinity for propofol and sevoflurane for intersubunit sites on GABA_A_ receptors. This result is consistent with a previous observation, using NMR, that a critical level of hydration was required for halothane to bind to proteins (65). To our knowledge, this is the first time the number of displaced water molecules has been correlated with affinity to explain GA specificity and selectivity.

In general, the proposed mechanism follows in a straightforward manner from the fundamental origin of the hydrophobic effect: water molecules able to hydrogen bond with only a few partners, as at an interface or within a cavity, can reduce their free energy by moving to bulk water. (Hydrophobicity is a slightly misleading term for this effect, since it is water that is “lipophobic” rather than the reverse). A broader study of ligand binding (66) identified a crucial role for water displacement in determining binding affinity to hydrophobic cavities, yielding the WaterMap docking corrections.

The number of displaced water molecules will increase with ligand-size and hydration of the cavity, suggesting that these factors will underlie affinity for the ligand and site respectively. This does not mandate that there is no role for hydrogen bonding between the general anesthetic and the protein in determining affinity. Cavities that are both hydrated and hydrophobic must balance the particular residue-residue interactions that favor the cavity with the surface tension of the water droplet that fills it. Polar side-chains can reduce that surface tension, especially for a small droplet.

In order to separate the role of dewetting from the role of hydrogen bonding between GA and sidechains, it was critical that we were able to compare binding to different homologous sites without varying the ligand. In principle our findings do not rule out a secondary contribution of hydrogen binding to polar side-chains to strengthening affinity: this would be enthalpically favorable, while still maintaining the entropic benefit of releasing cavity water. In practice, typical structure-activity experiments are unable to change ligand hydrogen bonding propensity without simultaneously introducing steric clashes, removing solubility, increasing pore block, and/or affecting dewetting. These conflicting contributions may explain the complex dependence observed by Harrison and colleagues for potency of a series of propofol analogs that potentiate GABA_A_ receptors, (67) or the fact that hydroxyl substitution with fluorine(38), but not bromine(67), removes general anesthetic function.

It is likely that the affinities we calculated may be particularly sensitive to the choice of initial structure and sampling of endpoint solvation. This could affect the overall predicted rank order of the four higher-affinity intersubunit propofol sites, and introduce the modest quantitative discrepancies with the results of Ref. (34). We would not expect these changes to affect the observed relationship between solvation and affinity. Consequently, our results suggest an intriguing possible origin for state-dependence of affinities: conformational changes that alter cavity solvation may directly affect ligand binding affinity for the cavity.

The second ongoing question we have addressed is the origin of the “orphan” site at the *α*^+^-*γ*^−^ interface. We propose here that the orphan site indirectly also reflects water displacement as a driver of affinity, because this lipid-accessible site has uniquely low hydration in the apo state. The *γ* M1 15′ contains an isoleucine residue (*γ* I242), which allows lipids to access the binding cavity, while the a and β subunits contain a methionine at the same location, which closes off the intersubunit site. This hypothesis could be tested experimentally by mutation of M1 15′ (I242M).

Overall, our results are consistent with results collected via mutagenesis and photolabeling involving GABA_A_ receptors over the past two decades, particularly those reported in Refs. (26, 29, 30, 34, 40, 58, 59, 67–70). These studies have focused exclusively on protein-mediated mechanisms of anesthetic binding. Nonetheless, membrane-mediated mechanisms underlying general anesthesia have remained intriguing to many researchers in part because of the compelling correlation of general anesthetic potency with lipophilicity first observed by Meyer and Overton (1), followed by numerous observations of the physical effects of anesthetics on lipid bilayers (71) at supertherapeutic concentrations. (72) Our results here suggest that this framing (lipid-mediated vs. protein-mediated) is misleading; general anesthetics don’t act <u>via</u> lipids, they act <u>like</u> lipids: amphipathic molecules that displace water from the protein surface and (in pLGICs) transmembrane cavities. This interpretation is consistent with the established sensitivity of the pLGIC family to lipids (73–78) and implies that lipid-specificity and general anesthetic specificity are deeply related questions.

## Author Contributions

SM: designed research; performed research; analyzed data; interpreted results; wrote the paper

JH: designed research; wrote the paper

GB: designed research; interpreted results; wrote the paper

## Acknowledgments

SM and GB were supported by research grant NIH P01GM55876-14A1. JH acknowledges support by the French government under grants DYNAMO (ANR-11-LABX-0011) and CACSICE (ANR-11-EQPX-0008). This project was supported with computational resources from the National Science Foundation XSEDE program through allocation NSF-MCB110149, as well as through the Rutgers Discovery Informatics Institute, which is supported by Rutgers and the State of New Jersey.

