## Supplemental Information for "Competitive dewetting underlies site-specific binding of general anesthetics to GABA(A) receptors"

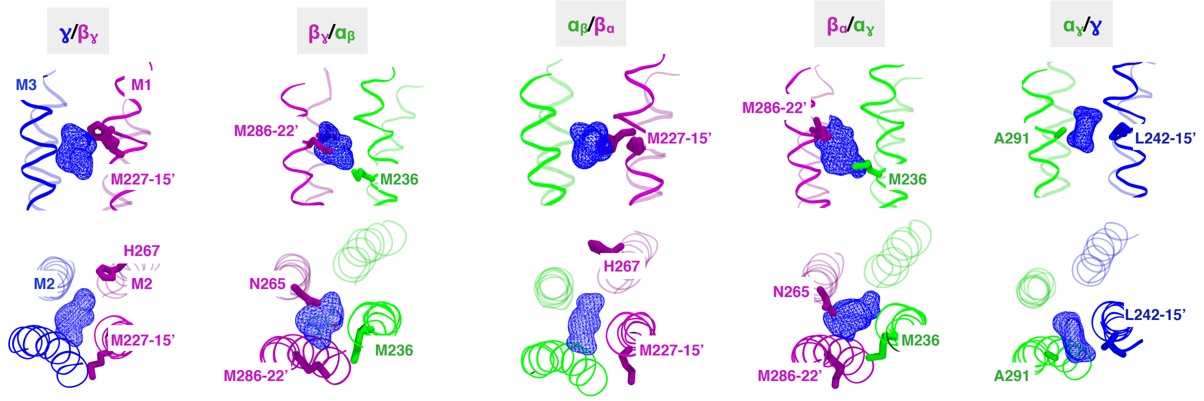


Figure S1: Individual subunit interfaces as labeled, with the protein colored by subunit, displaying the helices M3/M2 of a subunit and M1/M2 of another subunit, with view along TMD(top) and view from ECD(below). Licorice residues colored by subunit are the residues identified through previous experimental studies. Blue dots represent the average volume density of propofol over the course of entire simulation as calculated using the VOLMAP plugin of VMD ; $\alpha^+$-$\gamma^-$ site has no residues that have been experimentally identified and residues in licorice form in this site are residues homologous to ones shown in other sites.

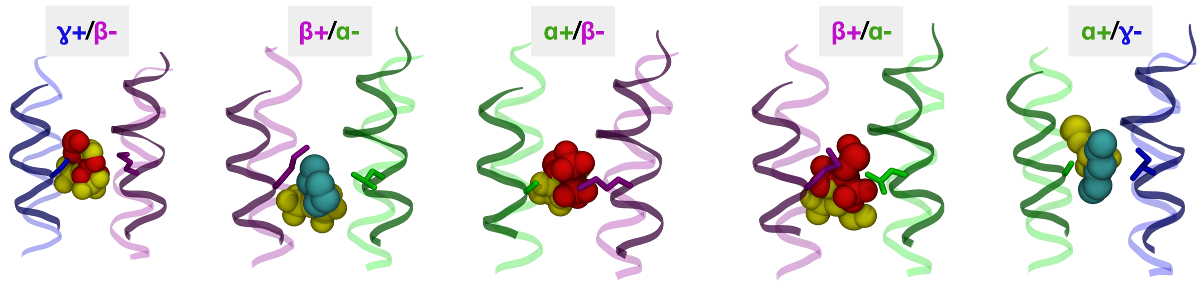


Figure S2: Different intersubunit sites viewed from the TMD, depicting the binding sites and orientation of sevoflurane/lipids identified through flooding simulation(sevoflurane-red; lipid-cyan) and standard MD simulation(yellow)

© 2013 The Authors
0006-3495/08/09/2624/12 $2.00 doi: 10.1529/biophysj.106.090944

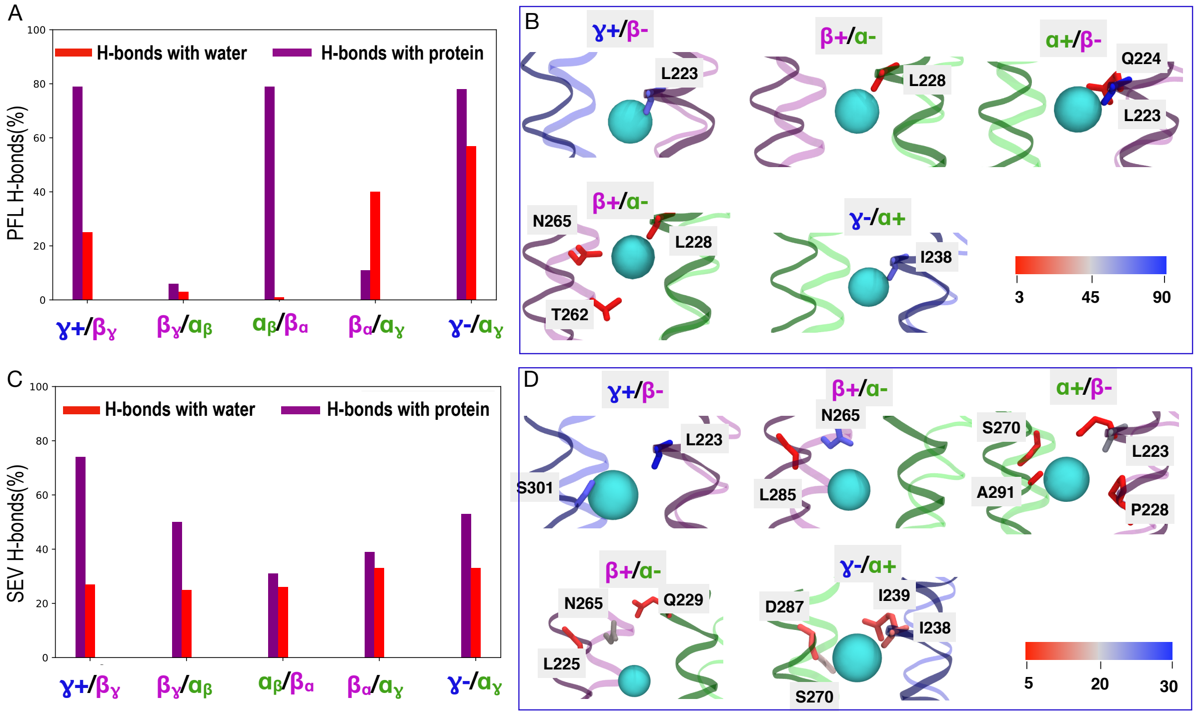


Figure S3: (A) Percentage of H-bonds between protein or water and (A) Propofol and (C) Sevoflurane. The protein residues that H-bond with PFL(B) and SEV(D) are shown in licorice and colored by the percentage of the hydrogen bonds formed in the course of the simulations, with red, denoting residues that forms least number of Hydrogen bonds with the ligand and blue, denoting the residues forming the highest number of hydrogen bonds.

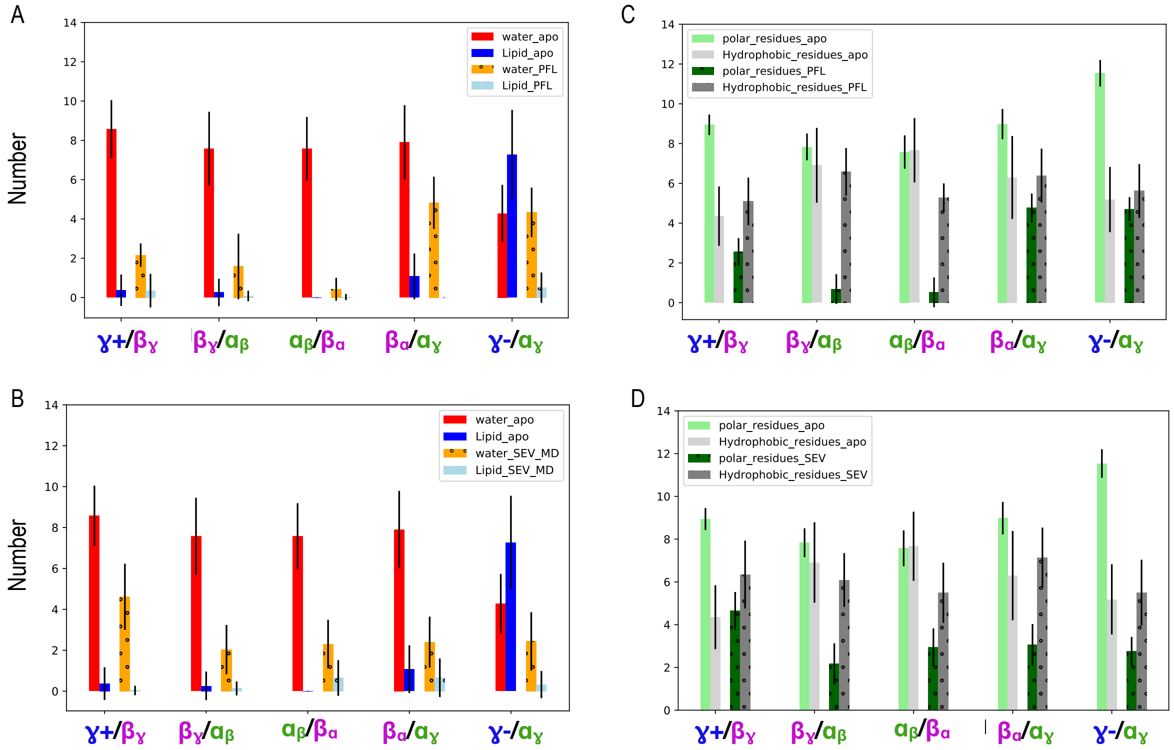


Figure S4: Comparisons of the number of water molecules and Lipid atoms at the different intersubunit site in Apo receptor system and propofol(A), sevoflurane (B) bound receptor system; Comparisons of the number of polar and hydrophobic protein present at the different intersubunit site in Apo receptor system and propofol(C), sevoflurane (D) bound receptor system.

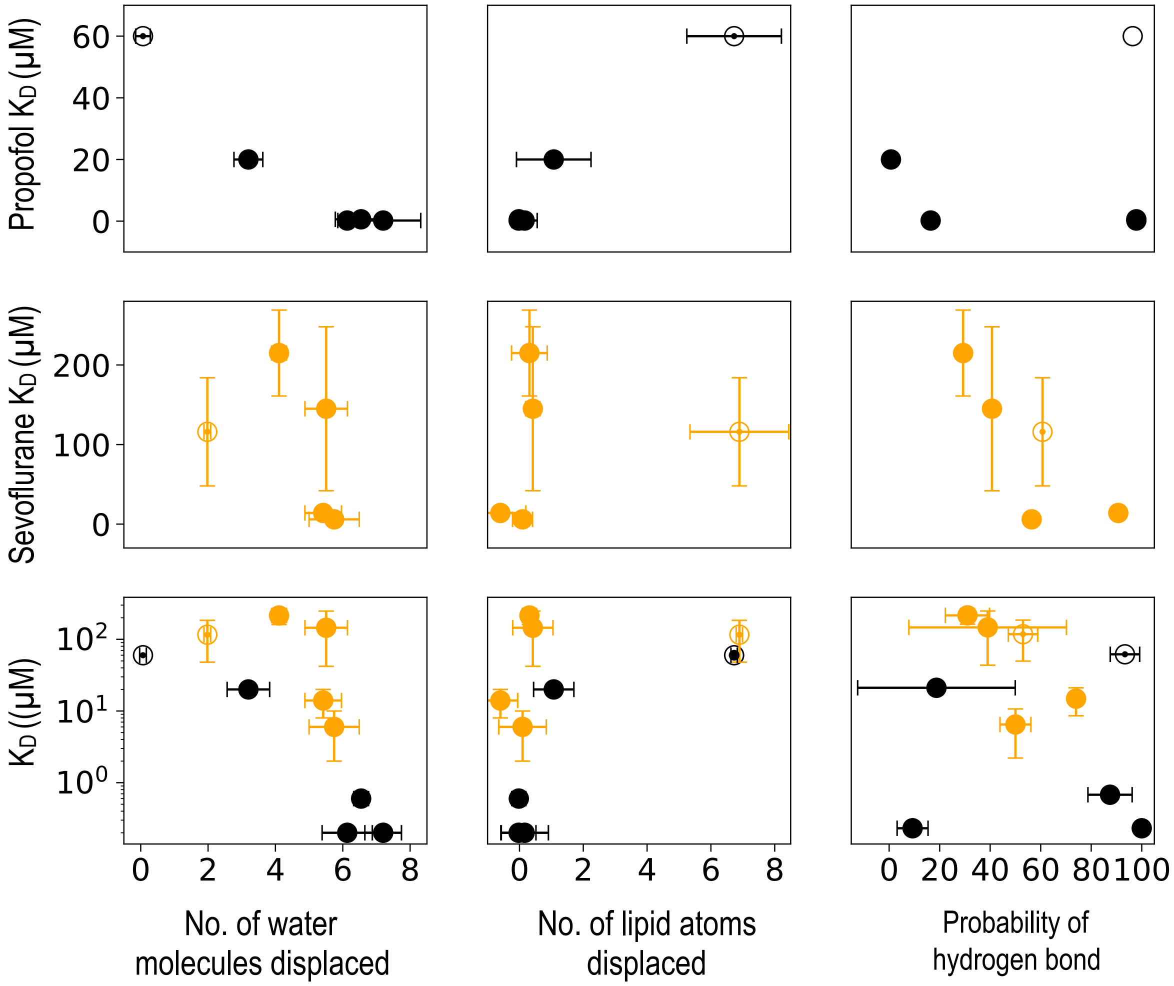


Figure S5: First and second row shows the correlation between binding site affinity and average no. of water and lipid atoms displaced and per- centage H-bond formation in the respective sites by propofol (black) and sevoflurane (orange) bound receptor system. Third row shows comparison of correlation between displacement of water and lipids by sevoflurane/propofol, and percentage H- bonding with their respective affinities on a Log scale. The open and closed circles represent sites that are termed as “lipid inaccessible” and “lipid accessible” respectively.
